# Dopamine D1 receptor signalling differentially regulates action sequences and turning behaviour in freely moving *Drosophila*

**DOI:** 10.1101/385674

**Authors:** Benjamin Kottler, Richard Faville, Jessika Bridi, Frank Hirth

**Affiliations:** King’s College London, Institute of Psychiatry, Psychology & Neuroscience, Maurice Wohl Clinical Neuroscience Institute, Department of Basic & Clinical Neuroscience, London, UK

**Keywords:** Action Selection, Decision Making, Central Complex, Dopamine, Drosophila

## Abstract

Here, we introduce a novel behavioural paradigm to study neural circuits and mechanisms underlying action selection and decision-making in freely moving ***Drosophila***. We first validate our approach by studying ***FoxP*** mutants and show that normally invariant patterns of motor activity and turning behaviour are altered in these flies, reminiscent of indecision. Then, focusing on central complex (CX) circuits known to integrate different sensory modalities and controlling premotor regions, we show that action sequences and turning behaviour are regulated by dopamine D1 (Dop1R1) receptor signalling. Dop1R1 inputs onto CX columnar wedge and ellipsoid body R2/R4m ring neuron circuits both negatively gate motor activity but inversely control turning behaviour. While flies deficient of D1 receptor signalling present normal turning behaviour despite decreased activity, restoring Dop1R1 level in R2/R4m-specific circuitry affects the temporal organisation of motor actions and turning. These findings suggest that columnar wedge and ring neuron circuits of the CX differentially modulate patterns of motor action sequences and turning behaviour by comparative Dop1R1 signalling for goal-directed locomotion.

## INTRODUCTION

Action selection is the process to “*do the right thing at the right time*” ^1^. It is a prerequisite for decision-making underlying goal directed locomotion, which requires the integration of sensory signals with internal states to translate them into action sequences. The most basic action selection for any ambulatory animal is between action and inaction, movement and immobility (Fig. 1a). Once activity is initiated, the sequence of activity bouts requires their correct organisation into action sequences. While speed can vary, moving from a starting point to a specific destination can be achieved by either one single bout or several successive bouts of activity. Moreover, the distribution of activity bouts can follow a scale-invariant pattern^2^. It has been shown that animals, from human to fly, have a tendency to initiate movement in bursts of activity rather than randomly. This is illustrated by the distribution of Inter-Bout Intervals (IBI) or pause between activity bouts which can follow a non-stochastic pattern, characterized by burstiness^2-4^. Goal-directed locomotion also includes decision-making processes that can be best summarized by “*you can’t turn left and right at the same time*”. For example when a moving animal is facing an obstacle, a decision is necessary to avoid collision ^5^ and a turn has to be executed (Fig. 1d). Since turning left and right at the same time is not possible, a selection between several available alternatives needs to be made regarding the nature of the turn by stopping and remaining inactive or changing direction by either turning left or right.

**Figure 1.**
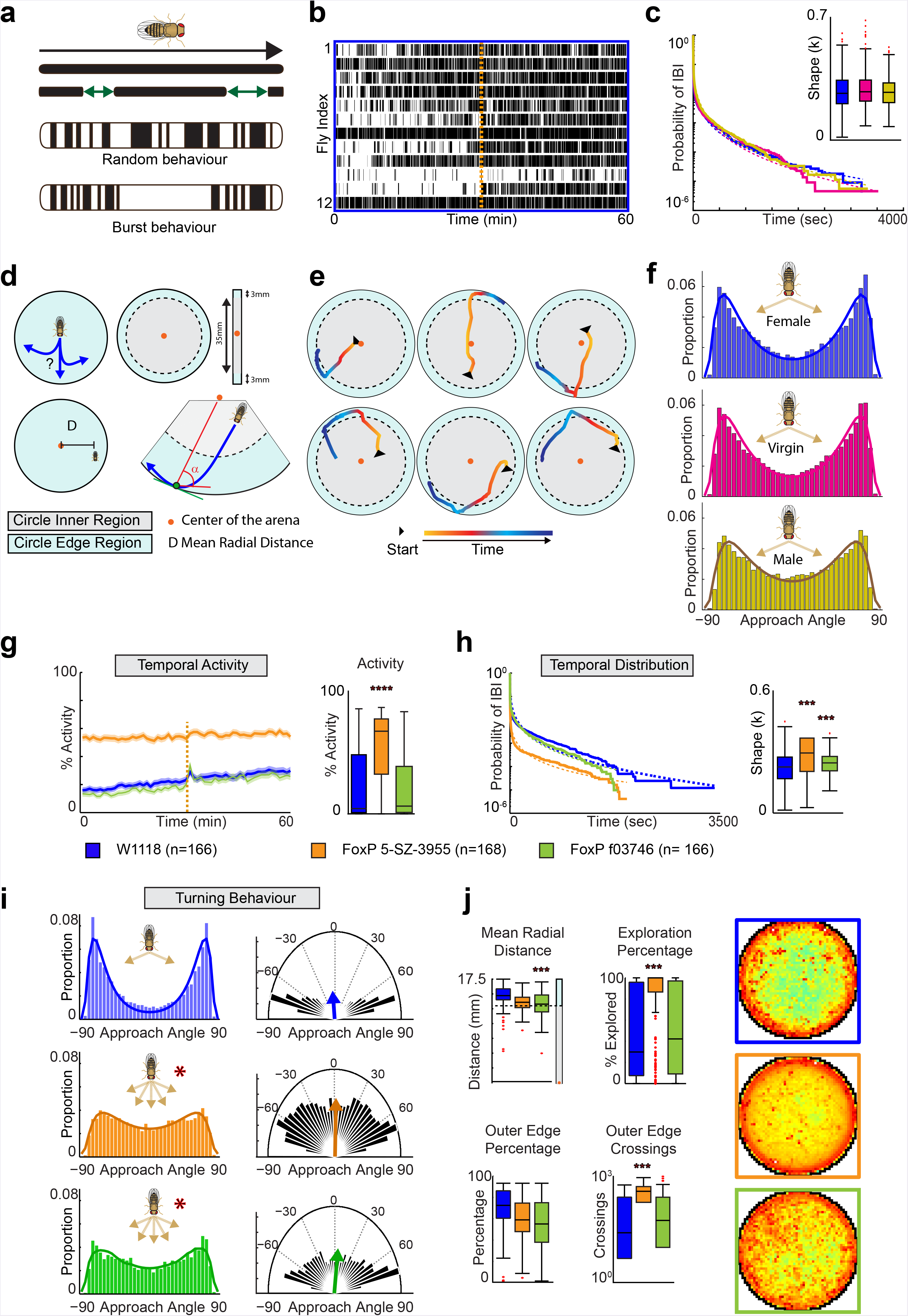
Invariant temporal distribution of motor activity and turning behaviour are affected in *FoxP* mutants. (**a**) Temporal sequence of activity bouts and inter-bout-intervals (IBIs) can follow a random distribution or burstiness where bouts of activity are clustered. (**b**) Raster plot of activity bouts (black bars) and IBIs (white) of 12 randomly chosen female flies; the dashed vertical orange line represents the mechanical stimuli applied after 30 min of recording. (**c**) Weibull plot of mean IBI distribution over time revealing a heavy tail; inset graph depicts shape factor κ which is a measure of random behaviour (=1) *vs* burstiness (<1); values are shown for three *w^1118^* control groups, females (blue), virgins (pink) and males (yellow). (**d**) Open-field arena divided into inner region (grey) and edge region (blue) delimited by 3mm distance from the edge; red dot represents arena centroid. The approach angle α is the deviation between the trajectory of the fly (in blue) when entering the edge region until contact with the wall (green dot) and the perpendicular at the point of contact to the centroid of the arena. (**e**) six examples of turning flies. (**f**) Approach angle distribution for *w^1118^* control groups, females, virgins and males, reveals bimodal distribution of turns with peaks at +/− 65-70 degrees. (**g-j**) Analysis of *FoxP^5 SZ-3955^* (orange) and *FoxP^03746^* (green) mutant flies, compared to *w^1118^* control (blue). (**g**) Temporal activity over 1 hour recording; dashed vertical line represents mechanical stimuli applied after 30 min of recording; right panel, mean activity over 60 min. (**h**) Temporal distribution of IBIs fitted to a Weibull distribution, and the dotted line showing the survival curve fitted response; right panel, shape factor κ. (**i**) Turning behaviour with approach angle distribution and polar plot. The fly cartoon represents bimodal distribution that is impaired in *FoxP* mutants. (**j**) Mean radial distance during turns (top left), mean exploration behaviour covering arena space (top middle), place preference shown as heat map (right), percentage spend in outer edge region (bottom left), and mean number of crossings between inner region and outer edge region (bottom middle). All data are mean +/−SEM; *p<0.5, **p<0.01, ***p<0.001.

In flies and other insects, several lines of evidence suggest that the central complex (CX) is involved in action selection and decision-making. The CX integrates various sensory cues and orchestrates motor output for adaptive behaviours which range from sleep, arousal and attention to higher motor control including goal-directed locomotion, orientation tuning, path integration and place learning^6-13^. The CX is a central brain structure composed of midline neuropils comprising the protocerebral bridge (PB), the fan-shaped body (FB), the ellipsoid body (EB), the noduli and the lateral accessory lobes (LAL). Two major types of projection neurons characterise the circuit architecture of the CX. Columnar-wedge neurons interconnect the different substructures of the CX, from PB to EB ^14-16^ and compartmentalize them into segments or modules, each of which corresponds to a segment of sensory space^7,17^. Tangential neurons form synaptic layers of the FB and EB. Previous studies identified the EB as a key node in mediating sensorimotor integration and action selection for goal-directed behaviour. These studies propose that visual cues and their position in space are represented in EB activity relative to the animal’s heading. Imaging ***in vivo*** studies revealed that a bump of activity restricted to a segment of the EB represents the angular orientation of an individual fly^18,19^. This suggests that columnar-wedge neuron activity encodes an internal compass and heading direction that combines visual landmarks with self-generated (idiothetic) cues^20,21^. Additionally, tangential ring neurons of the EB have been implicated in the regulation of visual place learning and visual orientation memory^22,23^. Together these data suggest that columnar and tangential ring neurons circuits converge onto the EB leading to the integration of self-generated cues and external sensory cues, two key features essential for correct navigation. However, despite detailed anatomical and physiological studies into CX functions, the neural mechanisms and molecular substrates mediating action selection and decision-making related to goal-directed locomotion are only starting to emerge.

Here, using a novel paradigm, we analysed the organisation of motor activity and turning behaviour in freely moving *Drosophila* and identified an invariant pattern of temporal activity and turning behaviour. We validated our approach by analysing well-described mutants of the forkhead box transcription factor gene, *FoxP* that are impaired in motor control and decision-making^24-26^. Analysis of their locomotion revealed that independent of activity levels, these *FoxP* mutants present variation in the distribution of IBIs and impaired turning behaviour. We then applied our novel paradigm to investigate the role of CX circuitry in action selection and decision-making. Targeted manipulations shown that columnar-wedge and tangential EB ring neuron circuits inversely regulate turning behaviour by D1 dopamine receptor Dop1R1 activity. Synaptically targeted *GFP reconstitution across synaptic partners* (*GRASP*) identified reciprocal connections between columnar-wedge and tangential ring neurons. Manipulation in R2/R4m but not other ring neuron circuitry of Dop1R1 levels in wild-type and in a *dumb2* heterozygote mutant background, revealed a specific role of the outer EB ring circuit for D1 receptor signalling. Our results demonstrate that CX circuit activity coordinates the temporal sequence of motor actions and turning behaviour, suggesting that comparative dopamine D1 receptor signalling between columnar wedge and EB R2/R4m neurons mediates their regulation.

## RESULTS

### Freely moving flies show invariant temporal patterns of activity and turning behaviour

To investigate motor action selection, we recorded the activity of freely moving flies in 35mm diameter arenas for 1 hour. Using the *Drosophila* ARousal Tracking DART system ^27^ (Supplementary Fig. 1a), we quantified motor action sequences and movement trajectories (Fig. 1b,e). Additionally, as a measure for the arousal state of a fly and a proxy for sensorimotor integration, we determined the response to sensory stimulation triggered by mechanical vibrations (Supplementary Fig. 1f). We first tested our system by recording the activity of *w^1118^* female, virgin and male control flies at 25°C and determined their temporal sequence of activity (respectively labelled in blue, pink and yellow in Fig. 1 and Supplementary Fig. 1). Fly motor activity can be represented as a raster plot where each bout of activity is indicated by a black box (Fig. 1b). Since animal activity is characterized by burstiness^2-4^, we also investigated the distribution of IBIs across the 60min window of recording. Similar to experiments that recorded activity over several days^4^, we found that the complementary cumulative distribution of IBIs fitted with the complementary cumulative Weibull distribution for all control groups (Fig. 1c). Weibull distributions are characterized by the scale parameter κ that reflects the degree of burstiness (Fig. 1c). A scale parameter κ of 1 corresponds to a random distribution (Poisson) of IBIs and its decrease corresponds to an increase in burstiness. For all control groups, we observed scale factors κ lower than 1 suggesting that even during 60min recording the temporal pattern of motor action sequences in *Drosophila* follows a non-stochastic distribution characterized by bursts of activity.

In parallel, we developed a paradigm to investigate turning behaviour as a proxy for decision-making. While freely moving in an open arena, flies tend to avoid the centre, a behaviour termed “centrophobism” ^28^. Despite their tendency to follow the wall, flies leave the edge to explore the central zone of the arena after which they return to the edge again. Thus, when re-approaching the edge of the arena, flies have to make a decision in order to avoid a collision with the wall, by achieving a turn that we can calculate. For that purpose, we divided the arena into two regions: the inner (centre) region and the edge (wall) region (Fig. 1d, light grey and light blue, respectively), with the edge region defined by a 3mm distance width which is at least twice as long as the body length of a fly. We then determined the mean radial distance for all control groups, which is the mean distance of a fly from the centre of the arena. We showed that their movement was circumscribed at the arena edge region. This allowed us to determine their place preference pictured with a heat map representation, which captures the percentage of time spent in the inner and outer region and the transitions between inner/centre to outer/edge region (Supplementary Fig. 1b). For the turning behaviour analysis, we specifically investigated turning events of flies transitioning from the inner region to the edge region within a continuous bout of activity (see methods for details). This restrictive definition allowed us to analyse necessary turns representing an inherent decision-making event even when perceptual choices were not obvious (Fig. 1e). We calculated the deviation angle between the fly’s trajectory and the perpendicular of the asymptote at the point of contact with the wall (denoted α in Fig. 1d and Supplementary Fig. 1c). These deviation angles could range from −90 and +90°, with a value of 0° representing a straight walk to the wall of the arena. Analysis of the proportional distribution of these approach angles for control flies revealed a bimodal distribution of turns with the most common modes or angles being around the absolute value of +/-70-80° (Fig. 1f). This bimodal distribution was extremely robust across gender, age, and time of day (Supplementary Fig. 1f). Together these results suggest that in addition to a non-stochastic temporal distribution of activity bouts, fly turning behaviour in a circular open arena follows an invariant spatial pattern characterized by a bimodal distribution.

### *FoxP* mediates motor action sequences and turning behaviour

In order to validate that our behavioural analysis can be used for studying action selection and decision-making, we first analysed *FoxP* mutants. *Drosophila FoxP* encodes a homologue of the *FoxP2* transcription factor which when mutated affects the temporal sequence of vocalisations in human, mice and songbirds^29,30^. Previous studies using the fruit fly revealed that *FoxP* mutants exhibit a defect in an operant self-learning but not in a world-learning assay^26^. This difference indicates that *FoxP* mutants are defective to assign value to self-generated actions but not for the assessment of external sensory cues. Previous studies also shown delayed decision-making in an odour discrimination task suggestive of indecision^24,31^. We used two previously characterized *FoxP* mutants, *FoxP^5SZ-3955^* and *FoxP^f03746^* and compared their behaviour with *w^1118^* control flies^24,26,31^. While the two *FoxP* mutants clearly presented different levels of activity (Fig. 1g and Supplementary Fig. 1d,e with *FoxP^5SZ3955^* and *FoxP^f0374β^* respectively in orange and green), they both exhibited a similar decrease in burstiness as illustrated by the increased shape factor κ compared to control (Fig 1h) indicative of defective action selection processing^4^.

Next we determined the turning behaviour of these *FoxP* mutants which identified a common behavioural difference compared to the control. While *w^1118^* flies shown a strong bimodal distribution of turns with peaks around +/- 70-80°, the two *FoxP* mutants revealed a more flattened distribution (Fig. 1i). These alterations in the distribution of the approach angle could not be attributed to differences in place preference as the 2D heat map still highlighted centrophobism (Fig. 1j). Moreover, the mean radial distances for both mutants were above the separation between inner and edge regions, even though *FoxP^f03746^* mutants were further away from the edges as depicted by the decrease in the radial distance (Fig. 1j). More importantly, neither of these two *FoxP* mutants shown any differences in the percentage of time spent in the outer edge region despite differences in both exploration and the absolute number of crossings for *FoxP^53955^,* which could be attributed to its higher activity level. In line with previous findings^24,26^, these data support a role for *FoxP* in motor control and decision-making and disentangle activity levels from burstiness and turning, suggesting that these behavioural manifestations are regulated by different mechanisms and/or neural circuits.

### Dopamine D1 receptor signalling in columnar wedge neurons modulates motor activity and turning behaviour

Modulatory dopamine receptor signalling is a key feature of action selection circuitries^9,32^. In *Drosophila,* Dop1R1 encodes a D1-like dopamine receptor that has been shown to be expressed in the CX^33,34^ and is required for a wide range of behavioural manifestations ranging from learning and memory to sleep and arousal ^35-38^. To study D1 function in our paradigm, we first examined whether its activity in columnar wedge neurons may regulate motor action sequences and turning behaviour. Activity of columnar wedge neurons has been shown to represent an internal compass and heading direction, which in the case of spatial navigation includes the execution of turning behaviour ^5,18,39^ We therefore hypothesized that targeted manipulation of D1 activity in columnar wedge neurons may impair motor action sequences and turning behaviour. To test this we used the previously^18,39^ characterized *R60D05-Gal4* driver specific to columnar wedge neurons (Supplementary Fig. 2a) and expressed *UAS-Dop1R1-IR* to impair D1 signalling. *R60D05>Dop1R1-IR* flies revealed significantly decreased activity levels but did not show alterations in burstiness. Interestingly, their approach angle distribution was statistically accentuated towards +/-75° angles, and thus more pronounced compared to controls (Fig. 2a). Detailed analysis of *R60D05>Dop1R1-IR* flies also revealed decreased centrophobism, even though they statistically explored less, probably due to the lowered activity level. In fact, they proportionally spent less time in the edge region, which might be indicating that their perception of the open arena could have been altered (Supplementary Fig. 2a).

**Figure 2.**
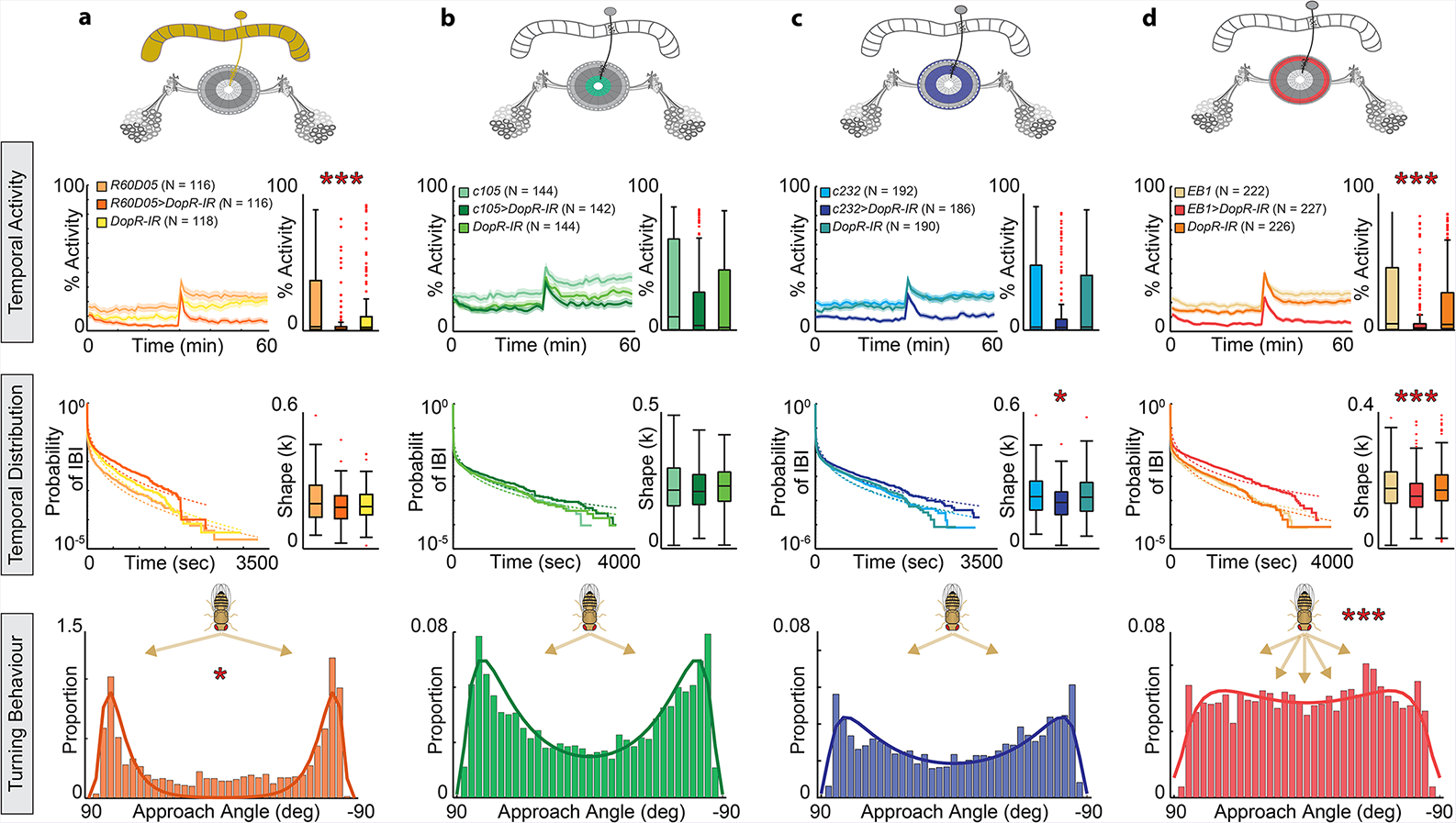
Columnar wedge and tangential ring neurons differentially modulate motor activity and turning behaviour via circuit-specific D1 receptor signalling. Colour-coded Gal4 expression pattern schematics for *UAS-DopR1-IR* expression by (**a**) *R60D05-Gal4* targeting columnar wedge neurons (orange), (**b**) *c105-Gal4* targeting ellipsoid body (EB) R1 circuitry (green), (**c**) *c232-Gal4* targeting R3/R4d circuitry (blue) and (**d**) *EB1-Gal4* targeting R2/R4m circuitry (red), and their respective controls *(Gal4/+* and *UAS/+*). Temporal activity resolution and activity percentage over the recorded hour; *p<0.5, **p<0.01, ***p<0.001.

### EB R2/R4m circuit-specific Dop1R1 function regulates temporal activity patterns and turning behaviour

We next studied tangential ring neurons and hypothesised that their dysfunction could affect the temporal pattern of motor actions and/or turning behaviour. To test this proposition, we used three Gal4 driver lines that have been widely used to characterize roles of EB R1-4 neurons in spatial orientation and place learning^19,23,40,41^. *c105-Gal4* shows expression in the inner ring layer (R1), *c232-Gal4* targets the middle (R3) and outer rim layer (R4d), and *EB1-Gal4* drives expression in the outer layers (R2 and R4m) (Supplementary Fig.S 2b-d). Thus, we examined the role of Dop1R1 by down regulating its expression in R1-4 ring neuron circuitries.

Expression of *UAS-DopR-IR* in R1 and R3/R4d layers, using c105 and c232 Gal4 driver lines respectively, did not modify activity levels (Fig. 2b,c). A slight but statistically not significant decrease could be observed with the *c232-Gal4* driver (Fig. 2c). This was accompanied with less exploration and the flies being significantly closer to the edge (Supplementary Fig. 2b,c). Interestingly, burstiness was altered in *c232- Gal4>DopR-IR* (Fig. 2c). Remarkably, driving *UAS-DopR-IR* with the R2/R4m driver line *EB1-Gal4* lead to a severe decrease in overall activity, which was accompanied with increased burstiness as revealed by the decreased *κ* factor compared to controls (Fig. 2d). Similar to R3/R4d-specific manipulations with *c232-Gal4, EB1-Gal4>DopR-IR* flies were more centrophobic and explored less the arena. Moreover, this R2/R4m specific overexpression of *UAS-DopR-IR* with the *EB1-Gal4* driver resulted in a change in the distribution of the angular approach, which no longer shown the typical bimodal distribution (Fig. 2d). These data identify that columnar wedge and EB R2/R4m neurons negatively gate motor activity, and that EB R2/R4m circuit-specific D1 signalling modulates the correct organisation of motor actions and turning behaviour.

### Columnar wedge neurons are reciprocally connected with tangential ring neurons

Given the striking opposite turning behaviour phenotypes mediated by columnar wedge and R2/R4m ring neurons, we wondered whether this might be due to functional interaction between these circuits. As already suggested by previous studies^5,18,39^, we postulated that columnar-wedge neurons would be connected with EB ring neuron circuitry and that ring neurons mediate motor action selection based on sensory representations encoded by columnar-wedge neuron activity.

In order to test this hypothesis, we used the *GFP reconstitution across synaptic partners (GRASP)* system to visualise potential synaptic connections between columnar-wedge and tangential ring neurons. GRASP utilizes the expression of two split-GFP halves that only fluoresce when reconstituted^42^. We applied a synaptically targeted GRASP system (syb-GRASP) and used *EB1-Gal4* specific to tangential R2/R4m ring neurons together with the *R60D05-LexA* driver specific to columnar wedge neurons (Supplementary Fig. 3).

We utilized split-GFP fragments *Aop-CD4::spGFP11* and *UAS-syb::spGFP1-10* that allow visualization of connections from ring to column, whereas a combination of *Aop-syb::spGFP1-10* and *UAS-CD4::spGFP11* allowed to visualize connections from column to ring. When combined, we detected GFP fluorescence around the circumference of the EB R2/R4m layer, restricted to predicted interactions of both singular expression patterns (Fig 3a, b). This GFP signal occurred in any combination indicative of potential reciprocal cell contacts between columnar wedge and ring neurons (Fig. 3c) as previously predicted by computational interrogation for goal-directed locomotion^5^ and CX circuit activity encoding heading direction^43,44^.

**Figure 3:**
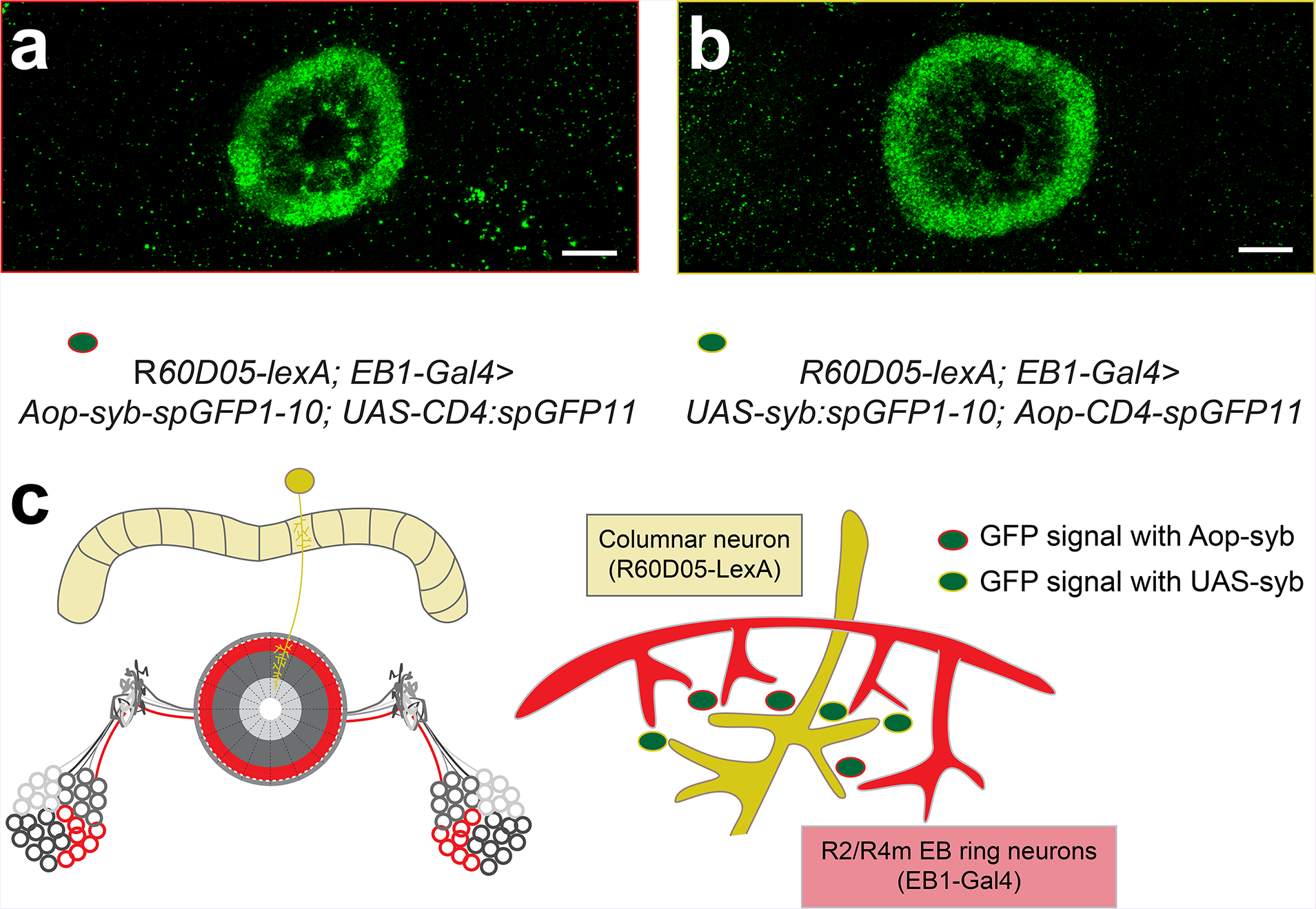
Synaptobrevin-tagged Green fluorescent protein *reconstitution across* columnar wedge and tangential ring neurons. (**a**) *R60D05-lexA; EB1-Gal4>Aop-syb-spGFP1 −10; UAS-CD4:spGFP11* visual reconstruction of GFP signal obtained with Aop-syb tagged GFP fragments (from columnar to tangential neurons). (**b**) *R60D05-lexA; EB1-Gal4>UAS-syb::spGFP1-10;Aop-CD4-spGFP11* indicate GFP signal obtained with UAS-syb tagged GFP fragments (from tangential to columnar neurons). (**c**) schematic of contact between columnar and R2.R4m tangential neurons on to the ellipsoid body (EB). Green dots with red frame indicate GFP signal obtained with Aop-syb tagged GFP fragments as in a; green dots with yellow frame indicate GFP signal obtained with UAS-syb tagged GFP fragments as in b. scale bar 20μm.

### Differential D1 receptor signalling in EB R2/R4m circuitry regulates motor action sequences and turning behaviour

In order to gain further insights into EB Dop1R1-mediated action selection and decision-making mechanisms, we investigated *Dop1R1* mutant flies. *Dumb^2^,* or *DopRf^02676^,* is a well-characterized strong hypomorphic allele^45^ that results in deficient learning and memory formation but also impairs arousal, sleep, as well as goal-directed and ethanol-induced locomotion^33,35-38^. Specifically, the *dumb^2^* mutation is caused by a P-element insertion in the first intronic region of the *DopR* gene^35^ resulting in a dramatic decrease of DopR protein levels^45^. This P-element contains a UAS cassette to which the Gal4 transcriptional activator can bind. By crossing the *Dumb^2^* mutant with a selective Gal4 driver, we can specifically restore Dop1R1 expression in neuronal subsets and circuits defined by the spatio-temporal activity of the respective Gal4 driver. In addition, this mutation does not show any compensatory effect on transcript levels for the DA receptors DopR2 and D2R2^36^.

We compared the *Dumb^2^* mutant with our control *w^1118^* flies. At our level of resolution and time scale of analysis, we observed a significant decrease of activity. *Dumb^2^* mutant flies shown an altered temporal pattern of activity with increased burstiness revealed by the decreased shape factor κ, confirming earlier findings that dopamine levels affect burstiness^4^. However, the turning behaviour of *Dumb^2^* mutant flies still revealed a bimodal distribution (Fig. 4a). Targeted manipulation of Dop1R1 in R1 or R3/4d layers with c105 and c232 driver lines, respectively, or their combination in *Dumb^2^* heterozygous background did not cause changes in any of the behavioural endpoints measured. Moreover, neither the pattern of burstiness nor the approach angle distribution, were affected (Supplementary Fig. S4a-b). In contrast, activity levels were affected by selectively targeting Dop1R1 level in the R2/R4m circuit with the *EB1-Gal4* driver in *Dumb^2^* heterozygous background. *EB1-Gal4/dumb^2^* flies were also characterised by increased burstiness illustrated by the decreased shape factor κ. Moreover, the bimodal distribution of turns was no longer detectable in *EB1-Gal4/dumb^2^* flies (Fig. 4b). These results pointed to a specific requirement of Dop1R1 signalling in R2/R4m circuitry. To further substantiate a specific requirement of Dop1R1 in this circuit, we selectively targeted Dop1R1 level in *Dumb^2^* heterozygotes background but also down regulated Dop1R1 level in wild-type background in R1, R3 and R4d together by combining *c105- Gal4* and *c232-Gal4* (Fig. 4c,d). However, neither the temporal organisation of motor actions, nor turning behaviour was affected in these flies. Together these results identify an EB R2/R4m circuit-specific requirement of Dop1R1 for action selection and decisionmaking in *Drosophila*.

**Figures 4:**
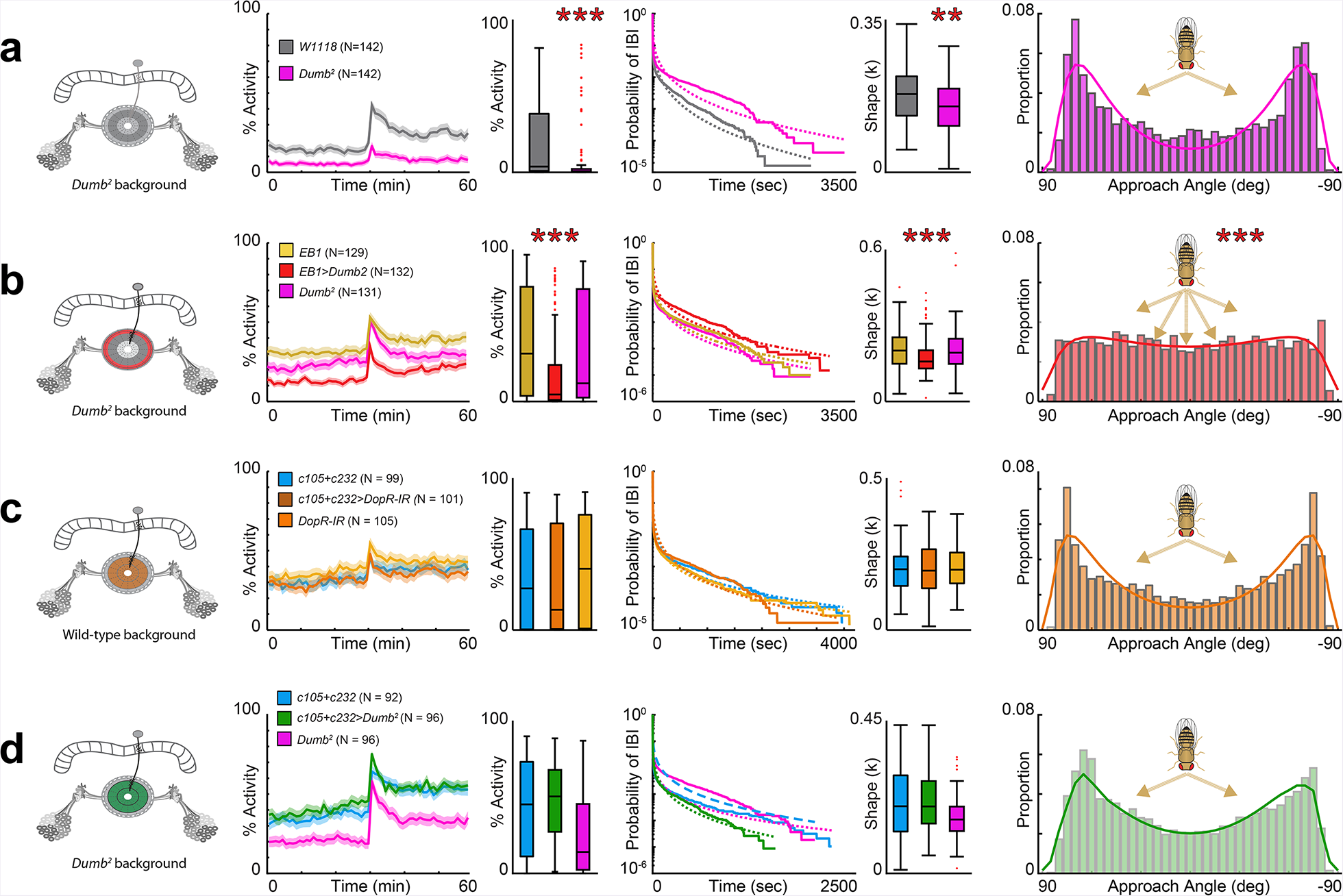
DopR1 receptor imbalance in ellipsoid body R2/R4m circuit affects temporal distribution of activity bouts and turning behaviour. Panels from left to right indicate DopR1 levels in entire brain and/or ring neuron subtype-specific ellipsoid body (EB) circuitry; temporal activity levels; mean activity; temporal distribution of inter-bout intervals (IBIs); shape factor κ; distribution of approach angle for (**a**) *Dumb^2^* mutant compared to *w^1118^* control flies; (**b**) R2/R4m specific *EB1-Gal4* targeting DopR1 in *Dumb^2^* heterozygous background. (**c**) R1+R3/4d specific *c105+c232-Gal4* expression of *UAS-DopR1-IR;* (**d**) R1+R3/4d specific *c105+c232-Gal4* targeting DopR1 in *Dumb^2^* heterozygous background. All data are mean +/-SEM; *p<0.5, **p<0.01, ***p<0.001.

## DISCUSSION

### Freely moving *Drosophila* present invariant patterns of activity and turning behaviour

Using a novel behavioural paradigm to investigate action selection and decisionmaking, we discovered invariant patterns of motor activity and turning behaviour in freely moving *Drosophila*. We observed a non-random distribution of IBIs that was comparable to recordings over several days^4^. The burstiness was a strong characteristic of every tested line, with a shape factor κ always <1 indicative of a none random distribution of IBIs (Fig. 1e). These results suggest that scale-invariant temporal pattern of locomotor activity follows a power law distribution seen in fractal behaviour^46^. Consistent with a previous report^46^, our findings therefore establish the EB as a central hub coordinating the temporal sequence of motor actions and burstiness, and identify specific dopamine D1 receptor signalling in columnar-wedge neurons and R2/R4m circuitry as key mediator for their regulation (fig. 2a, d).

Additionally, we observed a remarkably invariant pattern of turning behaviour when flies approached the arena wall. Our analysis revealed a bimodal distribution of approach angle with peaks of +/-70-80° irrespective of left and right preference (Fig. 1f). Since the turns we investigated occurred within a single uninterrupted bout of activity, the observed turning behaviour could revealed a decision-making process. But why do flies prefer to make a turn in such a way? We posit that flies know where they are in relation to the limits of the arena. As turning was measured within a single bout of activity, we hypothesise that flies anticipate the turn they are going to make in order to avoid a collision with the wall ^5^. Thus a +/-70-80° turn might represent the most efficient turning approach during a continuous walking bout, independently of activity levels as demonstrated by the *FoxP* mutant results (Fig. 1g-i).

### Impaired temporal distribution of motor activity and turning behaviour of *FoxP* mutants reminiscent of indecisiveness

Action selection and decision-making processes require the accumulation of perceptual evidences for adaptive behaviour, a process previously shown to be impaired in *FoxP* mutants^24,31^. Indeed, we observed altered motor activity and turning behaviour in two well-described *FoxP* mutants^24,26^, *FoxP^5SZ3955^* and *FoxP^f03746^*. Previous studies have shown these mutants to be deficient in a self-learning but not world-learning assay, suggesting that self-generated actions are not correctly processed or perceived^26^. These *FoxP* mutants also take more time to decide in an odour discrimination task indicative of decision-making deficits^24^. These behavioural alterations are accompanied by subtle anatomical alterations ^26^ but also by disrupted synaptic integration, affecting how salient information are added and retained ^31^. In our behavioural assay, these *FoxP* mutants showed decreased burstiness associated with increased shape factor κ, revealing an altered temporal distribution of IBIs. Such observations are indicative of an altered action selection process^2,4^. Additionally, we no longer detected the bimodal distribution of turns. As the measured turning events occurred within a continuous bout of activity, we assume that the difficulty of the task probably resides in the apparent absence of benefit whichever turn is completed and its fast execution, thus making a decision challenging. The changes in the angle approach distribution could be due to the fact that synaptic integration of self-generated and/or sensory cues is perturbed, as it has been shown for self-learning^26^ and odour discrimination tasks^31^. The resulting random distribution of turns in these *FoxP* mutants (Fig. 1f) suggests impaired action selection and the subsequently decision-making.

Despite these specific behavioural phenotypes, the expression and spatio-temporal activity of *FoxP* in *Drosophila melanogaster* remains unclear. There is no working antibody available and the expression patterns of different *FoxP-Gal4* driver highlight different structures. DasGupta and colleagues generated a *FoxP-Gal4* driver that shows restricted expression in the core of the Mushroom Body (MBs), the olfactory learning and memory centre^24^. However, Lawton and colleagues, generated another *FoxP-Gal4* driver by recombining the putative *FoxP* promoter with *UAS-CD8::GFP* for visualizing the expression pattern of FoxP protein. They found GFP-labelling in parts of the MBs and the entire protocerebral bridge (PB)^25^. The PB is involved in visual and tactile information processing^7^ necessary for spatial navigation and turning behaviour^5,18,39^. However it remains to be shown whether FoxP is indeed active in PB cells, especially the columnar wedge neurons.

### Columnar wedge and tangential ring neurons differentially modulate turning behaviour via circuit-specific D1 receptor signalling

Previous studies suggested that activity of columnar wedge neurons represent an internal compass that combines external sensory cues with self-generated cues^5,18,39^, thus exhibiting properties reminiscent of heading direction cells^21^. Based on these earlier findings, we first investigated the potential role of D1 signalling in columnar wedge neurons and found impaired activity levels and a bimodal distribution sharpened around peaks of +/- 75° turns. *R60D05-Gal>Dop1R1* mutant flies spent less time close to the wall of the arena (Fig. 2a and Supplementary 2a), indicative of altered place preference. What could be the role of D1 signalling for turning behaviour? Dopamine is a key neuromodulator known to be required for reinforcement signalling, thereby adding value to a behaviourally relevant signals^47,48^. In this sense, Dop1R1 signalling in columnar wedge neurons may code for a value with which the flies can evaluate their navigational goal in relation to their position in the arena and towards the wall. However, a possible limitation of our investigation is the *R60D05-Gal4* mediated expression of Dop1R1-IR in all columnar wedge neurons, which might be inappropriate to impact turning behaviour. Previous experiments by Green et al. (2017), using the same Gal4 driver line, revealed that a turn is performed when the activity of a single PB glomerulus takes over the rest of the population, leading to locally restricted neuronal activity. In fact, Green and colleagues shown that rotation of a tethered fly – equivalent to a turn in our assay - can be induced by local activation of 1-2 glomeruli only^39^. Thus, Dop1R1-mediated manipulation over the entire R60D05-targeted columnar neuron population might not have any consequence on the execution of turns. As we could not technically exploit this level of resolution with freely moving animals and investigate this hypothesis in our behavioural assay, we then extended our analysis to tangential ring neurons.

We first validated a computational assumption necessary for spatial navigation ^5^ and shown reciprocal contacts of potential pre-synaptic termini between columnar-wedge and ring neurons. Even though the physiology of the connections has to be experimentally demonstrated, the observed syb-GRASP related GFP reconstitutions support a model where columnar-wedge neurons, integrating visual landmark and heading direction^43^, and ring neurons, mediating an action selection process, together code for neural mechanisms underlying sensory integration and motor action selection for spatial navigation^5^. Therefore, we wondered how we could efficiently modulate this functional nexus in order to affect turning behaviour. More than changing activity levels of ring neurons, we decided to manipulate dopamine D1 signalling, a key modulator of action selection circuitries^9,32^, including the EB ^5,9^.

We found a profound effect of D1 signalling specifically in the EB R2/R4m circuitry. The temporal distribution of IBIs but also turning behaviour, were both affected. We observed an increase in burstiness and indecision when approaching the wall (Fig. 2d). Strikingly, the turning behaviour was not due to a general decrease of dopamine D1 receptor levels since *Dumb^2^* mutants were unaffected (Fig. 4a). Rather, the observed behavioural phenotypes were specifically caused by a Dop1R1 imbalance localized to the outer layer circuit of the EB, which our results suggest to be reciprocally connected with columnar-wedge neurons (Fig. 3). This behavioural observation was confirmed by targeting Dop1R1 in the R2/R4m circuitry in a heterozygous *Dumb^2^* mutant background, while manipulating R1-R3/4d EB circuits was without any measured behavioural consequence (Fig. 4). These findings are reminiscent of the inverted-U shape hypothesis of dopamine action and behavioural performance where both too little or too much dopamine signalling impairs behavioural outcome^49^. However, we extend this relation between dopamine signalling levels and behavioural performance to interconnected CX circuitry. In line with this concept, recent observations suggest that goal-directed locomotion requires the comparison of an internal heading direction signal with a navigational goal to guide turning choices ^43^. Together with these findings, these data propose that comparative dopamine D1 receptor signalling between columnar wedge and R2/R4m circuitry is modulating these adaptive behaviours.

## Author Contributions

Conceptualization B.K. and F.H.; Methodology B.K. and R.F.; Investigations B.K. and J.B.; Validation B.K.; Software R.F.; Writing - Original draft B.K.; Writing - Review & Editing B.K. and F.H.; Funding Acquisition F.H. Supervision B.K. and F.H.

## Acknowledgement

We thank Angelique Lamaze, Julien Colomb, Manolis Fanto and Bruno van Swinderen for comments on the manuscript and the Wohl Cellular Imaging Centre at King’s College London for help with microscopy. We are grateful to the Miesenbock lab for FoxP reagents and to Scott Waddel for Dopamine reagents. This work was supported by an IoPPN-King)s Independent Researcher Award to B.K., a PhD fellowship from CAPES Foundation-Ministry of Education of Brazil to J.B., and grants from the BBSRC (BB/N001230/1) and MRC UK (G0701498; MR/L010666/1) to F.H.

## METHODS

### Fly husbandry

Flies were maintained on a 12:12 light:dark cycle at 18°C and were grown standard cornmeal medium. Two days after eclosion, flies were anaesthesized with CO2, separated and kept at 25°C for at least 2 days until the experiment. Except for figure 1 which compares genders, all experiments used mated females.

### *Drosophila* Strains

The following strains were used: *W^118^* as control flies, *c105-Gal4, c232-Gal4, EB1-Gal4*. For GRASP experiments we combined *EB1-Gal4* and *R60D05-LexA* from the Janelia collection. The *Aop-CD4::spGFP11; UAS-syb::spGFP1-10* or *Aop-syb::spGFP1-10, UAS-CD4::spGFP11* were generous gifted from the Jepson lab. *Dumb^2^* line and *UAS-DopR-IR* (VDRC ID 107058) were generous gifts from Scott Waddell. *UAS-Rdl-IR* (VDRC ID 100429) was ordered from VDRC stock center. The FoxP, *FoxP ^5-SZ 3955^* and *FoxP^f03746^* mutants were generous gifts from Gero Miesenbock.

### Immunohistochemistry

Brains were dissected in cold PBS and fixed for 1hr at room temperature in 4% (wt/vol) paraformaldehyde in PBS (130 mM NaCl, 7 mM Na2HPO4,3 mM KH2PO4), followed by three 10-min washes in PBT (PBS containing 0.5% Triton X-100). Brains were then blocked in PBT plus 2% (wt/vol) NGS for 20-min, and incubated overnight at 4 °C with the primary antibodies diluted in the same solution at 4°C. The primary antibodies were mouse anti-GFP (ThermoFisher Scientific 33-2600, 1:500 for msGFP detection or Sigma-Aldrich G6539, 1:200 for reconstituted splitGFP detection). The secondary antibodies were goat anti-mouse and anti-rabbit conjugated to Alexa fluor 488 or 555 (Invitrogen Molecular Probes, 1:150). For *R60D05-Gal4>UAS-mCD8::GFP* brains, no anti-GFP antibody was used. This was followed by three 10- min washes in PBT, 2-h incubation with the secondary antibodies at room temperature, three 20-min washes in PBT again, and one final wash in PBS. Tissues were mounted in Vectashield (Vector Laboratories). Images were acquired with a Nikon A1R confocal microscope equipped with 40x 1.3NA Plan Fluor oil immersion objective, processed using the Fiji software and figures constructed in Adobe Illustrator.

### Open-field behavioural analysis

Fly tracking was performed using custom-made platforms made of Acetal copolymer (POM-C), Tecaform AH. A platform comprises 36 open-field arenas, each arena 35mm in diameter and 1.5mm height, all covered with a transparent acrylic sheet with 1mm holes for breathing. The platform with arenas was placed on a white light plate that provided uniform cold light illumination within a temperature-controlled incubator (Stuart Scientific). Only females where taken unless mentioned otherwise. Flies were transferred individually into each arena after a short cold anesthesia and left to acclimatize for at least 30 minutes. Video-assisted motion tracking and analyses were carried out using the DART system ^27^ with custom-made MATLAB (Mathworks) scripts. Video recordings were carried out using a Logitech c920 camera at 10 frames per second for 1 hour. The positions of flies were extracted every 2 recorded frames (0.20 sec).

### Kinematic calculations

To determine locomotion parameters, we calculated fly motion as follows: The inter-frame displacement, *D_i_* (which is calculated between the *i^th^* and (*i*-1)^th^ video frames) is calculated using the following equation:

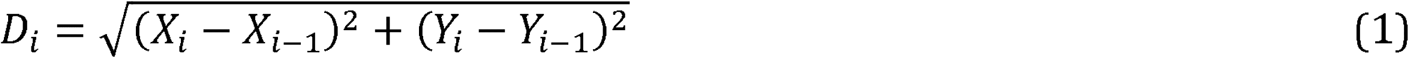

where *X_J_/Y_j_* denote the *X*/*Y* coordinates (respectively) for frame *j*. The inter-frame speed is calculated using Eqn. (1) as follows:

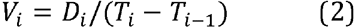

where *T_j_* denotes the time stamp for frame *j*. The activity threshold of a fly: a fly is considered “active” for a given video frame if Eqn. (2) exceeds 2mm/sec, which is equivalent to the travelling the body length of the fly every second.

### Motor actions

The *percentage activity* is calculated as the percentage of video frames during which the fly is active. This data is also shown in a *raster plot* that depicts activity bouts during 60 minutes of recording, with black vertical lines representing active frames and white space representing pauses as inter-bout intervals (IBIs). *Traces:* plots showing the track of each individual fly during the first 10 minutes of recording within the corresponding genotypic group. *Activity/min:* activity scores for each animal were calculated and these were combined with all other recorded flies per genotype to obtain a mean *activity/min* for all flies of that genotype. *Speed* was calculated by averaging Eqn. (2) for each video frame over all flies per genotype. *Distance* was calculated by summing Eqn. (1) over all active frames for each fly (during the 60 minutes of recording), and then averaged over all recorded flies per genotype.

### Motor action sequences

*Action initiation* was defined as the number of times a fly transitions from an inactive to active state divided by the total time spent inactive. This metric is averaged over all recorded flies per genotype to give the mean number of activity bouts started per second. *Mean bout length* was defined as the duration of an activity bouts during which an individual fly was continuously active. Measures are averaged over all recorded flies per genotype. *Interbout interval* (IBI) is the time between two bouts of activity when the fly is continuously inactive; the minimum IBI is 0.2 seconds, which corresponds to a single video frame. This metric is averaged over all recorded flies per genotype to give the mean duration of an IBI as pause.

### Burstiness behaviour

Animal activity is characterized by burstiness, which characterises burst of activity in a short time window followed by extensive periods of inactivity (Barabasi, 2005; Reynolds, 2011; Sorribes et al., 2011). Burstiness can be described by a Weibull distribution, which calculates the cumulative distribution of inter-bout intervals (IBIs) of a given group of animals (here a genotype) during a defined period of time, here during the 60min window of recording. To quantify the IBI distribution, survival curves were fit using a Weibull distribution with the following functional form:

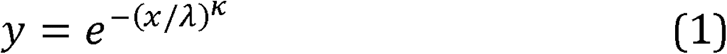

where *x* is the IBI, *λ* is the shape factor, and *κ* the scale factor. Using this Weibull distribution, *λ* and *κ* can be deduced that reflects the degree of burstiness for a given group. A shape factor *κ* of 1 corresponds to a random distribution (Poisson) of IBIs whereas its decrease corresponds to an increase in burstiness behaviour, that is: burst of activity are no longer randomly distributed but occur in clusters.

In order to estimate the parameters *λ* and *κ* Eqn. (1) is linearised by using the transform *y̅* = *log_e_(~log_e_(y)*) and *x̅ = log_e_(x)* such that:

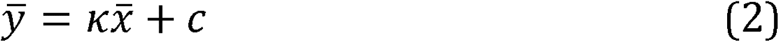

where *c* = *−κlog_e_*(*λ*). From Eqn. (2), the parameters *λ* and κwere estimated using the Matlab optimization function, *fit* (Mathworks).

### Mechanical stimulus response and arousal state

A mechanical stimulation was delivered after 30 minutes of recording and was composed of 5 vibrations at 3 Volts for 200ms separated by 800ms. Mechanical stimulation and response analyses were carried out as previously described ^27^. The pre-stimuli speed represents the average speed of all flies per genotype 2 minutes prior to mechanical stimulation (calculated using Eqn. (2)). The absolute post-stimuli speed is calculated again using Eqn. (2), but is smoothed (so as to remove high-frequency noise) using the following equation:

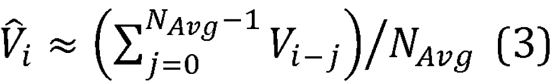

where *N_Avg_* is the number preceding video frames (*N_Avg_* = 10). The final absolute post-stimuli speed trace is determined by interpolating Eqn. (3) to the nearest second. From the post-stimuli speed trace, the amplitude is calculated as the difference between the maximum stimuli response and the average pre-stimuli speed. To quantify the features of the stimuli response, the relative post-stimuli signal (calculated as the absolute post-stimuli speed minus the average pre-stimuli speed) is fitted with a single-inactivation exponential equation with the following functional form:

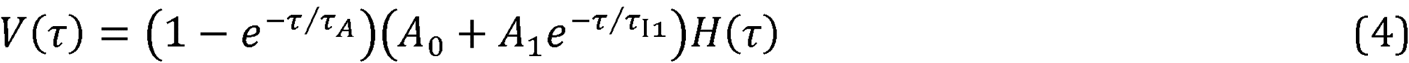

where *τ = t − δt, H*(*t*) is the Heaviside step function (=1 if *t >* otherwise 0),*A*_0-2_are scale factors *τ*_*A*_ is the activation time constant, and *τ_A_/τ_B_* are inactivation time constants. The exponential equation parameters (the scale factors, time constants and *δt*) for (4) were fitted using the Matlab optimization function, *fit* (Mathworks).

### Turning Behaviour Metrics

In order to extract the distribution of approach angles, several parameters are considered:

- **Movement Threshold ≤ 2mm/sec**. The speed at which a fly is considered moving.
- **Circumferential Distance = 2mm**. The circumferential threshold distance for determining movement after edge contact. The circumferential distance is the ditancwe the fly travels around the circular region (as opposed to radially)
- **Pre-edge Contact Duration = 2 sec**. The active duration before contact with the wall used to determine the approach angle.
- **Post-edge Contact Duration = 2 sec**. The active duration after contact with the wall used to determine edge behaviour.
- **Minimum Edge Contact = 1 sec**. Minimum duration the fly has to be in the outer ring to be considered as edge region.
- **Outside Edge Distance = 3mm**. The distance from outside edge whereby the fly is considered in contact with wall.

### Radial Distance and Circumferential angle

A fly position has the following coordinates at a given time T_(i)_ over an experiment of n_Frm_ Frames: (X_(i)_, Y_(i)_). The radial distance and circumferential angle are denoted, respectively, by *ρ_(i)_* =*D*(*X_(i)_*,*Y_(i)_*) and *φ_c(i)_* =ω(*X_(i)_*,*Y_(i)_*) where:

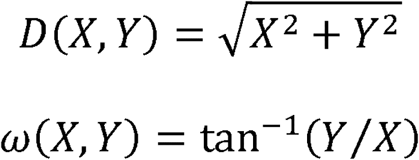

### Edge Region

We define the “edge region” of the circle to be the annulus formed by the circle’s edge and an inner circle of radius (R-ΔR). The distance R-(R-ΔR) is manually established within the DART program at 3mm.

**Figure 1.**
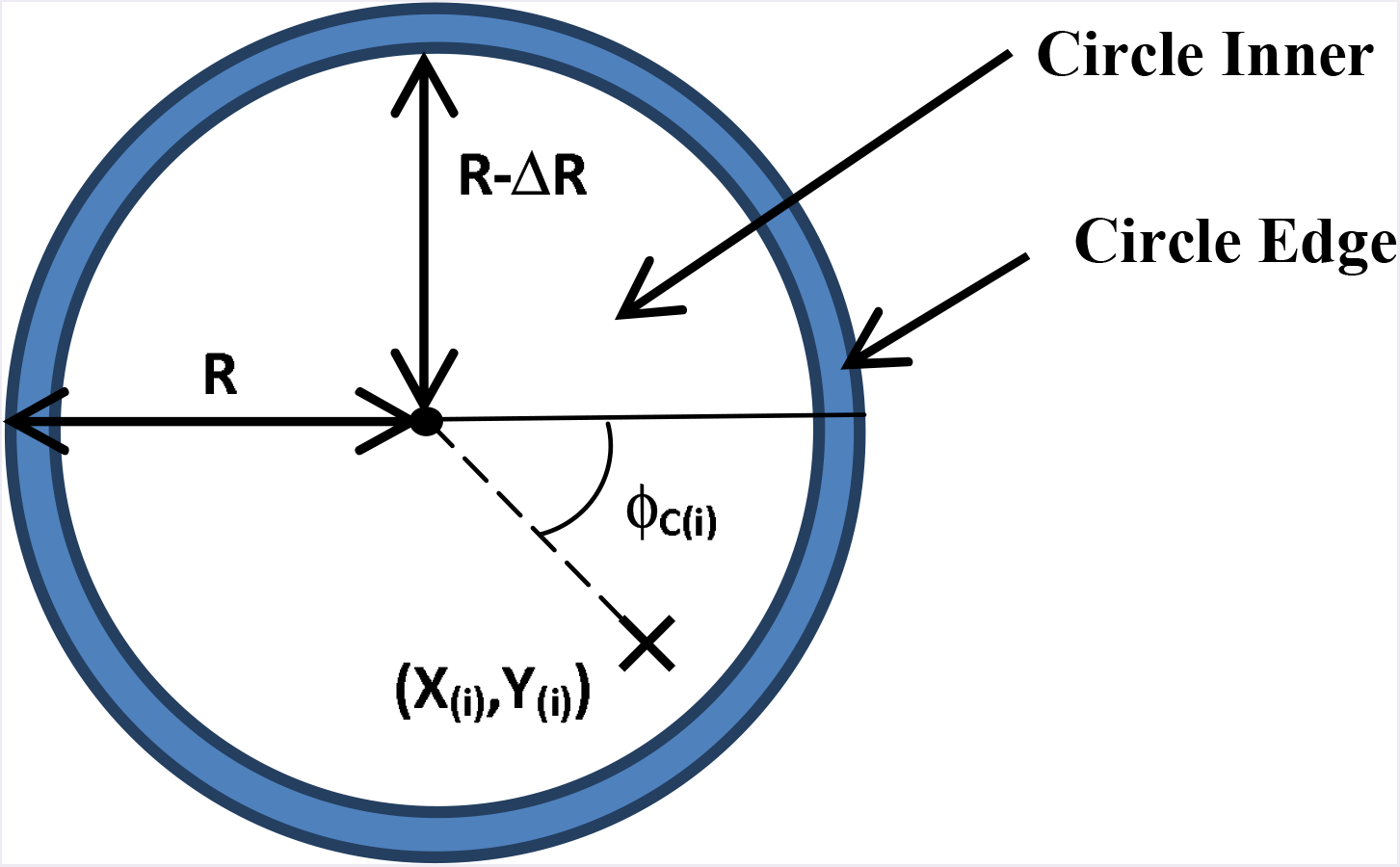
Schematic diagram illustrating regions and other definitions.

### Edge Contact

A fly is considered to be within the circle’s edge region if the following is true:

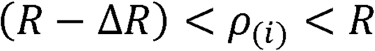

Given this is true, a valid edge contact event is said to have occurred if the fly remains in the outer region for at least T_m_in, or in terms of video frames:

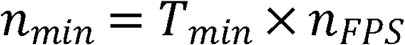

where ***n**_FPS_* is video frame rate and equals 5 frame/sec. From this, video frames are given a designation of 1 if that frame constitutes a valid edge contact (0 otherwise) and is denoted by the symbol, *Z*_(*i*)_. It follows that the proportional duration that the fly spends in the outer region, denoted by *P_out_*, is calculated as follows:

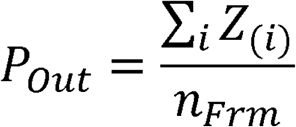

### Outer Region Crossings

Given the valid edge contact events are known, it is trivial to determine the times at which the fly enters/leaves the outer region (after a valid contact event) which has cardinality, respectively. Therefore, the total number of edge crossings is given by:

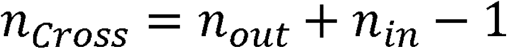

### Pre/Post-Edge Contact Sequences

In order to calculate the approach angle the pre-edge contact subsequence is formed by taking the n_B_ frames preceding ξ_(*j*)_. Conversely, the post-edge contact behavioural metrics are determined from the subsequences formed from the n_A_ frames subsequent to ξ_(*j*)_ The values of n_B_/n_A_ are linked to the video frame rate by the equation:

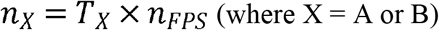

where T_B_/T_A_ is the pre/post-contact duration, respectively. Note that for an edge contact to be considered valid for the metric calculations, the following must be true:

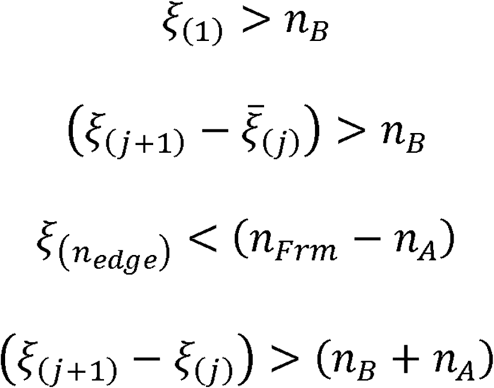

In other words, there must be at least A) n_B_ frames before the start of the first edge contact event, B) n_B_ frames between a fly leaving and re-entering the edge region, C) n_A_ frames subsequent to the final edge contact event, and D) (n_A_ + +n_B_) frames between the start of contact events.

### Approach Angle

The positional coordinates of the pre-edge contact subsequences are rotated with respect to *φ_ξ(j)_)* (i.e., the circumferential angle from the frame where the fly first crosses into the edge contact region):

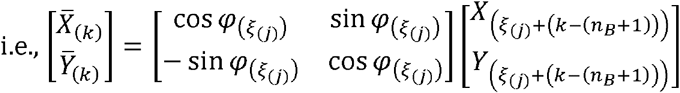

where k = 1 to n_B_ = 1

The approach angle (denoted by φ_A(j)_) is calculated from the rotated coordinates by taking the distance weighted sum of the inter-frame angles as follows:

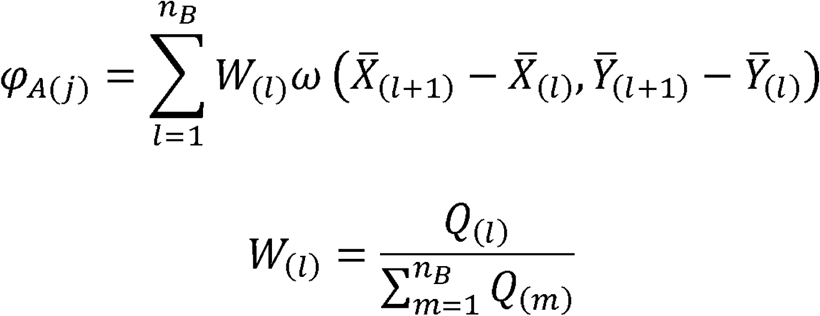

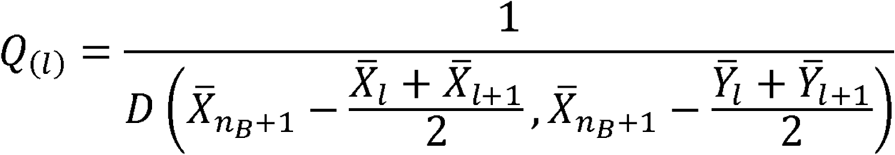

In other words, the inter-frame angles are weighted by calculating the distance from the average position between adjacent frames to the final frame in the sub-sequence (i.e., the frame where the fly enters the edge region). This means that inter-frame angles that are closer to the edge region have a higher weighting. Negative approach angles denote that the fly is approaching the edge from the left side, and positive angles approaching from the right.

Note that only the frames where the fly is continuously moving toward the arena edge are considered for the approach angle calculation 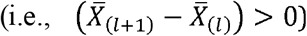. Furthermore, for the metric analysis, the approach angles are separated into bins of size Δφ from a domain of −90^°^ to 90^°^. In our analysis, we choose a resolution of 5degrees. This leads to plotting histogram value (Y value) in function of orientation bin angle (X value).

For the polar plot graphical representation of the turning preference, each histogram is normalized to the maximum Y-value. This way the longest value reaches the outside of the circle and all other values are relative to this maximum and we can plot a vector length representing the average turning event.

### Overall Fly Location

This is the ratio of the frames where the fly is within the edge region to the total number of frames:

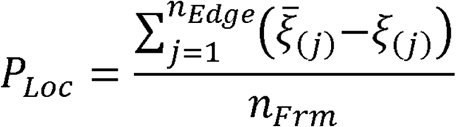

### Exploration Percentage

To determine the percentage of the open-field region that the fly explores during the duration of the experiment, the positional coordinates are converted from mm to pixels as follows:

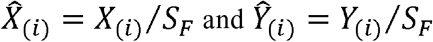

Where *S_F_* is the pixel to mm scale factor, which for our experiments, was usually on the order of 0.35 mm/pixel. From these converted positional coordinates, the locations of the fly between each frame is estimated by linear interpolation and rounded to the nearest pixel (integer) value. Given the number of unique locations, from the known and interpolated positional values which is denoted by *n_Tot_* the exploration percentage is given by the equation:

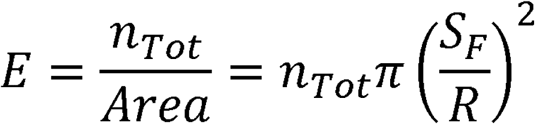

### Statistical analysis

For the histograms, each data set was tested for normality using the Anderson-Darling test with α = 0.05. If every data set under comparison was normal and the variances were similar (a ratio of 4 between the highest and lowest variance was used as the cut-off), then a one-way ANOVA test was used to determine whether any differences existed between groups. If significance was found for ANOVA with α = 0.05, then pair-wise comparisons were carried out using a post hoc Tukey-Kramer test, again with α = 0.05. If any of the data sets was found not to be normally distributed, then a Kruskal-Wallis test was used to determine any overall differences between the groups with α = 0.05. If significance was achieved, a post hoc pair-wise Mann-Whitney U test with Dunn-Sidak correction was used to compare groups with α = 0.05. For each test group, two controls were used. All calculations were performed using MATLAB within the DART software. For testing the significance of changes in the bimodal distribution of approach angles, we first pooled together turns of similar absolute values, assuming that conceptually a right or a left turn is the same. We then performed two sample Kolmogorov-Smirnov (KS) tests on the normalized data to the maximum between the experimental and the two parental controls to evaluate alteration in the distribution of approach angles. Significance was achieved when the experimental was statistically different from the two controls, and that no difference was observed between controls. All data were extracted from the DART software and KS tests were run in Graph Pad 7.04. In all figures, the p-value of the statistical test is represented as either one star (p<0.05), two stars (p<0.01) or three stars (p<0.001).

## Data availability

Data supporting the findings of this study are available within the paper and its Supplementary Information files and from the authors upon request.

## REFERENCES

1 Prescott, T. J., Bryson, J.J. & Seth, A. K. Introduction. Modelling natural action selection. Philosophical transactions of the Royal Society of London. Series B, Biological sciences 362, 1521–1529, doi: 10.1098/rstb.2007.2050 (2007).

2 Barabasi, A. L. The origin of bursts and heavy tails in human dynamics. Nature 435, 207–211, doi:10.1038/nature03459 (2005).

3 Reynolds, A. M. On the origin of bursts and heavy tails in animal dynamics. Physica A 390, 245–249, doi: 10.1016/j.physa.2010.09.020 (2011).

4 Sorribes, A., Armendariz, B. G., Lopez-Pigozzi, D., Murga, C. & de Polavieja, G. G. The origin of behavioral bursts in decision-making circuitry. PLoS Comput Biol 7, e1002075, doi: 10.1371/journal.pcbi. 1002075 (2011).

5 Fiore, V. G., Kottler, B., Gu, X. & Hirth, F. In silico Interrogation of Insect Central Complex Suggests Computational Roles for the Ellipsoid Body in Spatial Navigation. Frontiers in behavioral neuroscience 11, 142, doi:10.3389/fnbeh.2017.00142 (2017).

6 de Bivort, B. L. & van Swinderen, B. Evidence for selective attention in the insect brain. Current opinion in insect science 15, 9–15, doi:10.1016/j.cois.2016.02.007 (2016).

7 Heinze, S. Unraveling the neural basis of insect navigation. Current opinion in insect science 24, 58–67, doi:10.1016/j.cois.2017.09.001 (2017).

8 Pfeiffer, K. & Homberg, U. Organization and functional roles of the central complex in the insect brain. Annual review of entomology 59, 165–184, doi:10.1146/annurev-ento-011613- 162031 (2014).

9 Strausfeld, N. J. & Hirth, F. Deep homology of arthropod central complex and vertebrate basal ganglia. Science 340, 157–161, doi:10.1126/science.1231828 (2013).

10 Strauss, R. The central complex and the genetic dissection of locomotor behaviour. Current opinion in neurobiology 12, 633–638 (2002).

11 Turner-Evans, D. B. & Jayaraman, V. The insect central complex. Current biology : CB 26, R453–457, doi:10.1016/j.cub.2016.04.006 (2016).

12 Varga, A. G., Kathman, N. D., Martin, J. P., Guo, P. & Ritzmann, R. E. Spatial Navigation and the Central Complex: Sensory Acquisition, Orientation, and Motor Control. Frontiers in behavioral neuroscience 11, 4, doi:10.3389/fnbeh.2017.00004 (2017).

13 Donlea, J. M. Neuronal and molecular mechanisms of sleep homeostasis. Current opinion in insect science 24, 51–57, doi:10.1016/j.cois.2017.09.008 (2017).

14 Ito, K. & Awasaki, T. Clonal unit architecture of the adult fly brain. Adv Exp Med Biol 628, 137–158, doi:10.1007/978-0-387-78261-4_9 (2008).

15 Lin, C. Y.et al. A comprehensive wiring diagram of the protocerebral bridge for visual information processing in the Drosophila brain. Cell reports 3, 1739–1753, doi: 10.1016/j.celrep.2013.04.022 (2013).

16 Wolff, T., Iyer, N. A. & Rubin, G. M. Neuroarchitecture and neuroanatomy of the Drosophila central complex: A GAL4-based dissection of protocerebral bridge neurons and circuits. The Journal of comparative neurology 523, 997–1037, doi:10.1002/cne.23705 (2015).

17 Strausfeld, N. J. Arthropod Brains: Evolution, Functional Elegance and Historical Significance. (Cambridge MA: Harvard University Press). (2012).

18 Seelig, J. D. & Jayaraman, V. Neural dynamics for landmark orientation and angular path integration. Nature 521,186–191, doi:10.1038/nature14446 (2015).

19 Seelig, J. D. & Jayaraman, V. Feature detection and orientation tuning in the Drosophila central complex. Nature 503, 262–266, doi:10.1038/nature12601 (2013).

20 Green, J. & Maimon, G. Building a heading signal from anatomically defined neuron types in the Drosophila central complex. Current opinion in neurobiology 52, 156–164, doi:10.1016/j.conb.2018.06.010 (2018).

21 Heinze, S. Neuroethology: unweaving the senses of direction. Current biology: CB 25, R1034–R1037, doi:10.1016/j.cub.2015.09.003 (2015).

22 Kuntz, S., Poeck, B., Sokolowski, M. B. & Strauss, R. The visual orientation memory of Drosophila requires Foraging (PKG) upstream of Ignorant (RSK2) in ring neurons of the central complex. Learning& memory 19, 337–340, doi:10.1101/lm.026369.112 (2012).

23 Ofstad, T. A., Zuker, C. S. & Reiser, M. B. Visual place learning in Drosophila melanogaster. Nature 474, 204–207, doi:10.1038/nature10131 (2011).

24 DasGupta, S., Ferreira, C. H. & Miesenbock, G. FoxP influences the speed and accuracy of a perceptual decision in Drosophila. Science 344, 901–904, doi:10.1126/science. 1252114 (2014).

25 Lawton, K. J., Wassmer, T. L. & Deitcher, D. L. Conserved role of Drosophila melanogaster FoxP in motor coordination and courtship song. Behav Brain Res 268, 213–221, doi: 10.1016/j.bbr.2014.04.009 (2014).

26 Mendoza, E.et al. Drosophila FoxP mutants are deficient in operant self-learning. PloS one 9, e100648, doi:10.1371/journal.pone.0100648 (2014).

27 Faville, R., Kottler, B., Goodhill, G. J., Shaw, P. J. & van Swinderen, B. How deeply does your mutant sleep? Probing arousal to better understand sleep defects in Drosophila. Scientific reports 5, 8454, doi:10.1038/srep08454 (2015).

28 Besson, M. & Martin, J. R. Centrophobism/thigmotaxis, a new role for the mushroom bodies in Drosophila. Journal of neurobiology 62, 386–396, doi:10.1002/neu.20111 (2005).

29 Fisher, S. E. & Scharff, C. FOXP2 as a molecular window into speech and language. Trends Genet 25,166–177, doi:10.1016/j.tig.2009.03.002 (2009).

30 Haesler, S.et al. Incomplete and inaccurate vocal imitation after knockdown of FoxP2 in songbird basal ganglia nucleus Area X. PLoS biology 5, e321, doi:10.1371/journal.pbio.0050321 (2007).

31 Groschner, L. N., Chan Wah Hak, L., Bogacz, R., DasGupta, S. & Miesenböck, G. Dendritic Integration of Sensory Evidence in Perceptual Decision-Making. Cell, doi:10.1016/j.cell.2018.03.075 (2018).

32 Cisek, P. & Kalaska, J. F. Neural mechanisms for interacting with a world full of action choices. Annual review of neuroscience 33, 269–298, doi: 10.1146/annurev.neuro.051508.135409 (2010).

33 Alekseyenko, O. V., Chan, Y. B., Li, R. & Kravitz, E. A. Single dopaminergic neurons that modulate aggression in Drosophila. Proceedings of the National Academy of Sciences of the United States of America 110, 6151–6156, doi:10.1073/pnas.1303446110 (2013).

34 Kim, Y. C., Lee, H. G., Seong, C. S. & Han, K. A. Expression of a D1 dopamine receptor dDAl/DmDOPl in the central nervous system of Drosophila melanogaster. Gene Expr Patterns 3, 237–245 (2003).

35 Kim, Y. C., Lee, H. G. & Han, K. A. D-l dopamine receptor dDAl is required in the mushroom body neurons for aversive and appetitive learning in Drosophila. Journal of Neuroscience 27, 7640–7647, doi:10.1523/Jneurosci.1167-07.2007 (2007).

36 Kong, E. C.et al. A pair of dopamine neurons target the D1-like dopamine receptor DopR in the central complex to promote ethanol-stimulated locomotion in Drosophila. PloS one 5, e9954, doi:10.1371/journal.pone.0009954 (2010).

37 Riemensperger, T.et al. Behavioral consequences of dopamine deficiency in the Drosophila central nervous system. Proceedings of the National Academy of Sciences of the United States of America 108, 834–839, doi:10.1073/pnas.1010930108 (2011).

38 Ueno, T.et al. Identification of a dopamine pathway that regulates sleep and arousal in Drosophila. Nature neuroscience 15,1516–1523, doi:10.1038/nn.3238 (2012).

39 Green, J.et al. A neural circuit architecture for angular integration in Drosophila. Nature 546, 101–106, doi:10.1038/nature22343 (2017).

40 Neuser, K., Triphan, T., Mronz, M., Poeck, B. & Strauss, R. Analysis of a spatial orientation memory in Drosophila. Nature 453, 1244–1247, doi:10.1038/nature07003 (2008).

41 Pan, Y. F.et al. Differential roles of the fan-shaped body and the ellipsoid body in Drosophila visual pattern memory. Learning& memory 16, 289–295, doi:10.1101/lm. 1331809 (2009).

42 Macpherson, L. J.et al. Dynamic labelling of neural connections in multiple colours by trans-synaptic fluorescence complementation. Nature communications 6, 10024, doi:10.1038/ncomms10024 (2015).

43 Green, J., Vijayan, V., Mussells Pires, P., Adachi, A. & Maimon, G. Walking Drosophila aim to maintain a neural heading estimate at an internal goal angle, doi:10.1101/315796 (2018).

44 Kim, S. S., Rouault, H., Druckmann, S. & Jayaraman, V. Ring attractor dynamics in the Drosophila central brain. Science 356, 849–853, doi:10.1126/science.aal4835 (2017).

45 Lebestky, T.et al. Two different forms of arousal in Drosophila are oppositely regulated by the dopamine D1 receptor ortholog DopR via distinct neural circuits. Neuron 64, 522–536, doi:10.1016/j.neuron.2009.09.031 (2009).

46 Martin, J., Faure, P. & Ernst, R. The power law distribution for walking-time intervals correlates with the ellipsoid-body in Drosophila. J Neurogenet 15, 205–219 (2001).

47 Schultz, W., Stauffer, W. R. & Lak, A. The phasic dopamine signal maturing: from reward via behavioural activation to formal economic utility. Current opinion in neurobiology 43, 139–148, doi:10.1016/j.conb.2017.03.013 (2017).

48 Waddell, S. Reinforcement signalling in Drosophila; dopamine does it all after all. Current opinion in neurobiology 23, 324–329, doi:10.1016/j.conb.2013.01.005 (2013).

49 Birman, S. Arousal mechanisms: speedy flies don)t sleep at night. Current biology : CB 15, R511–513, doi: 10.1016/j.cub.2005.06.032 (2005).

